# A myristoyl switch at the plasma membrane triggers cleavage and oligomerization of Mason-Pfizer monkey virus matrix protein

**DOI:** 10.1101/2023.12.20.572496

**Authors:** Markéta Častorálová, Jakub Sýs, Jan Prchal, Anna Pavlů, Lucie Prokopová, Zina Briki, Martin Hubálek, Tomáš Ruml

**Author notes:** Contributed equally (Markéta Častorálová, Jakub Sýs, Jan Prchal).

## Abstract

For most retroviruses, including HIV, association with the plasma membrane (PM) promotes the assembly of immature particles, which occurs simultaneously with budding and maturation. In these viruses, maturation is initiated by oligomerization of polyprotein precursors. In contrast, several retroviruses, such as Mason-Pfizer monkey virus (M-PMV), assemble in the cytoplasm into immature particles that are transported across the PM. Therefore, protease activation and specific cleavage must not occur until the preassembled particle interacts with the PM. This interaction is triggered by a bipartite signal consisting of a cluster of basic residues in the matrix (MA) domain of Gag polyprotein and a myristoyl moiety N-terminally attached to MA. Here, we provide evidence that myristoyl exposure from the MA core and its insertion into the PM occurs in M-PMV. By a combination of experimental methods, we show that this results in a structural change at the C-terminus of MA allowing efficient cleavage of MA from the downstream region of Gag. This suggests that, in addition to the known effect of the myristoyl switch of HIV-1 MA on the multimerization state of Gag and particle assembly, the myristoyl switch may have a regulatory role in initiating sequential cleavage of M-PMV Gag in immature particles.

## Introduction

Interaction with the plasma membrane (PM), budding, and maturation of viral particles are the final steps in retroviral replication cycles. In most retroviruses, the Gag-PM interaction is mediated by a bipartite motif consisting of a highly basic region and the N-terminal myristoyl (myr) moiety of MA. Despite similarities in the main mechanisms of viral capsid assembly and membrane targeting, there are remarkable morphogenetic differences between retroviruses. One significant difference lies in the location of the assembly sites; these are located at the PM for C-type retroviruses such as HIV and in the cytoplasm for D-type retroviruses, such as Mason-Pfizer monkey virus (M-PMV). The preassembled intracytoplasmic viral particles (ICAPs) of D-type retroviruses are then transported to the PM for budding (Rhee & Hunter, 1990).

Retroviral MA proteins are the key players in targeting Gag to the PM. HIV-1 MA is myristoylated and exists in two conformational states dictated by its location. In the cytoplasm, the myristoyl (myr) moiety is sequestered inside the hydrophobic pocket of MA. Upon Gag interaction with the PM, myr is expelled from the MA molecule and inserted into the lipid bilayer by the so-called myr switch (Tang et al., 2004). In HIV-1, the myr exposure is triggered by the interaction of the MA domain of Gag with phosphatidylinositol-4,5-bisphosphate [PI(4,5)P_2_], which is present exclusively in the PM (Ono et al., 2004; Saad et al., 2006; Tang et al., 2004). The equilibrium between exposed and sequestered myr states in HIV-1 MA is concentration and oligomerization-dependent, in contrast to myristoylated HIV-2 MA, which is exclusively monomeric *in vitro*. This can be explained by the tighter sequestration of the myr chain within the protein core (Saad et al., 2008). In D-type retrovirus M-PMV, MA myristoylation is also essential for the targeting of immature particles to the PM (Rhee & Hunter, 1991), but with a significantly lower affinity for water-soluble PI(4,5)P_2_ compared to HIV-1 MA. In addition, the interaction with this PM component fails to induce the myristoyl switch in purified M-PMV (myr+)MA *in vitro* (Kroupa et al., 2016; Prchal et al., 2012).

After the interaction of Gag or pre-assembled viral particles with the PM and budding, maturation occurs. The proteolytic maturation involves a delicately regulated sequential processing of polyprotein precursors (Konvalinka et al., 2015). The importance of sequential cleavage of HIV-1 Gag to liberate the mature structural proteins matrix (MA), capsid (CA), nucleocapsid (NC), and p6 and the spacer peptides SP1 and SP2 has been recognized for decades (Ericksonviitanen et al., 1989; Pettit et al., 1994; Tritch et al., 1991). However, our understanding of how this coordinated event is regulated remains incomplete.

Recently, the challenge of analyzing a population of viruses in heterogeneous life cycle phases was overcome with an approach using a photo-destructible HIV protease inhibitor (Pettit et al., 2005). This allowed synchronization of the immature virus particles and triggering of polyprotein processing with light. However, it also uncoupled maturation from assembly, preventing study of the interplay between these processes and the impact of Gag-membrane interactions on proteolytic cleavage. Recently, experiments using nanoscale flow cytometry and instant structured illumination microscopy demonstrated that activation of HIV-1 protease occurs during viral assembly prior to release of the virus (Tabler et al., 2022). However, fluorescence lifetime imaging microscopy and single-virus tracking revealed that there is a delay between HIV-1 particle assembly and maturation (Qian et al., 2022). The importance of precise regulation of maturation was demonstrated by overexpression of HIV-1 protease which, in addition to its cytotoxic effects, prevented HIV-1 particle formation and budding (Karacostas et al., 1993). Similar effect was also shown for Rous sarcoma virus (RSV) where premature proteolytic processing also lead to budding defects (Xiang et al., 1997).

*In vitro* data suggested that initial cleavage of HIV-1 Gag occurs at the SP1/NC site, followed by cleavage at the SP2/p6 and MA/CA sites, and finally at the NC/SP2 and CA/SP1 sites (Pettit et al., 1994; Wiegers et al., 1998; Wondrak et al., 1993). Researchers observed a similar pattern of ordered processing in infected cells by using mutants in which individual cleavage sites were disrupted by point mutations (Wiegers et al., 1998). These data led to the conclusion that the cleavage events at individual sites occur with different kinetics. The initial fast cleavage at the C terminus of SP1, releasing NC, allows condensation of the ribonucleoprotein core. Next, CA is liberated from the MA, and finally, the release of SP1 from CA initiates capsid condensation and core formation. Interestingly, even a small proportion of uncleaved Gag has a dominant negative effect on HIV-1 infectivity, as shown by Müller *et al*. (Muller et al., 2009). Blocking cleavage of MA/CA site had the most pronounced impact, exhibiting a transdominant negative effect (Lee et al., 2009). Both MA-CA and MA-CA-SP1 maturation products accumulated in cells when the MA/CA cleavage site was blocked by point mutations (de Marco et al., 2010; Mattei et al., 2018; Muller et al., 2009). Similarly, in murine leukemia virus (MLV), partially cleaved Gag interferes with infectivity when present at equimolar concentrations with the mature proteins (Rulli et al., 2006).

In M-PMV, the protease domain of Gag-Pro and Gag-Pro-Pol polyproteins can theoretically dimerize in ICAPs, however, processing is not initiated until the particles reach the PM (Parker & Hunter, 2001). Thus, the maturation of D-type retroviruses must be tightly regulated and the mere Pro dimerization is probably not sufficient for effective ICAP maturation. *In vitro* analysis of the proteolytic processing of intracytoplasmic M-PMV particles also suggested a sequential manner of Gag maturation. However, the first protein domain released by M-PMV protease from Gag was MA, with a 10-fold greater affinity for the MA/PP cleavage site than other Gag cleavage sites (Pichová et al., 1998). Parker and Hunter observed similar cleavage kinetics for M-PMV MA/PP and PP/p12 cleavage sites in COS-1 cells (Parker & Hunter, 2001). This is in contrast with data on HIV-1, for which initial cleavage occurs at SP1/NC (Wiegers et al., 1998), and RSV, for which *in vivo* studies showed that the MA-p2 junction is the most slowly cleaved site (Xiang et al., 1997). This was supported by *in vitro* cleavage of peptides mimicking the cleavage sites in RSV Gag, in which both the MA-p2 and p2-p10 junctions were cleaved very inefficiently compared to the other sites (Cameron et al., 1992).

Interestingly, the HIV-1 (myr+)MA V7R mutant, in which myr is sequestered in the MA core, was shown to be defective in Gag processing in HeLa and 293T cells (Hikichi et al., 2019; Ono & Freed, 1999). This was confirmed in CD4(+) T cells, which are the natural target cells of HIV-1 (Lee et al., 1998). In M-PMV, the possible role of MA myristoylation in maturation was also suggested (Prchal et al., 2011). Even though M-PMV MAPP (MA C-terminally extended with part of the downstream phosphoprotein [PP] domain of M-PMV Gag) can be cleaved efficiently by M-PMV protease *in vitro*, the N-terminally myristoylated MAPP [(myr+)MAPP] is cleaved very poorly (Prchal et al., 2011). Further, the NMR solution structure of myristoylated MAPP showed a short alpha helix spanning the position of the MA/PP cleavage site that is closely associated with the myristoyl moiety (Prchal et al., 2012).

Data presented in this study indicate that the interaction of M-PMV Gag with the PM can significantly enhance both trimerization and cleavage of MA from the rest of Gag, suggesting it may serve as an additional mechanism for maturation control. More broadly, we demonstrate that the interaction of the N-terminal myristoyl with the PM can be projected to the C-terminal region of MA to increase the availability of the domain for downstream processes such as proteolytic cleavage.

## Results

### Interaction of M-PMV (myr+)MAPP with liposomes triggers MA myristoyl switch and subsequent proteolytic processing at the MA/PP junction

M-PMV maturation occurs upon the interaction of M-PMV ICAPs with the PM during and after budding. However, as we have already shown, the myristoylated M-PMV MA extended with 18 amino acid residues from the downstream PP domain of Gag and a His-tag anchor [ (myr+)MAPP] is cleaved very poorly by M-PMV protease (Prchal et al., 2011). The mechanism that could trigger proteolytic processing of membrane-bound myristoylated Gag is the exposure of myristoyl from the hydrophobic pocket of MA, the myristoyl switch. Therefore, we sought to confirm that interaction with liposomes enables cleavage at the MA/PP junction in (myr+)MAPP. We compared the efficiency of cleavage of both M-PMV (myr-)MAPP and (myr+)MAPP bound to liposomes mimicking the phospholipid composition of the PM inner leaflet (Doktorova et al., 2017). As previously observed (Prchal et al., 2011), in the absence of liposomes, (myr-)MAPP was quickly cleaved by M-PMV protease (**Fig. 1A**), but cleavage of (myr+)MAPP was significantly delayed (**Fig. 1B**). Our new data reveal that binding of (myr+)MAPP to liposomes significantly enhances the cleavage kinetics (**Fig. 1B**). Therefore, we propose that the interaction of (myr+)MAPP with liposomes triggers the myristoyl switch that subsequently enables efficient cleavage at the MA/PP junction of M-PMV (myr+)Gag.

**Fig. 1.**
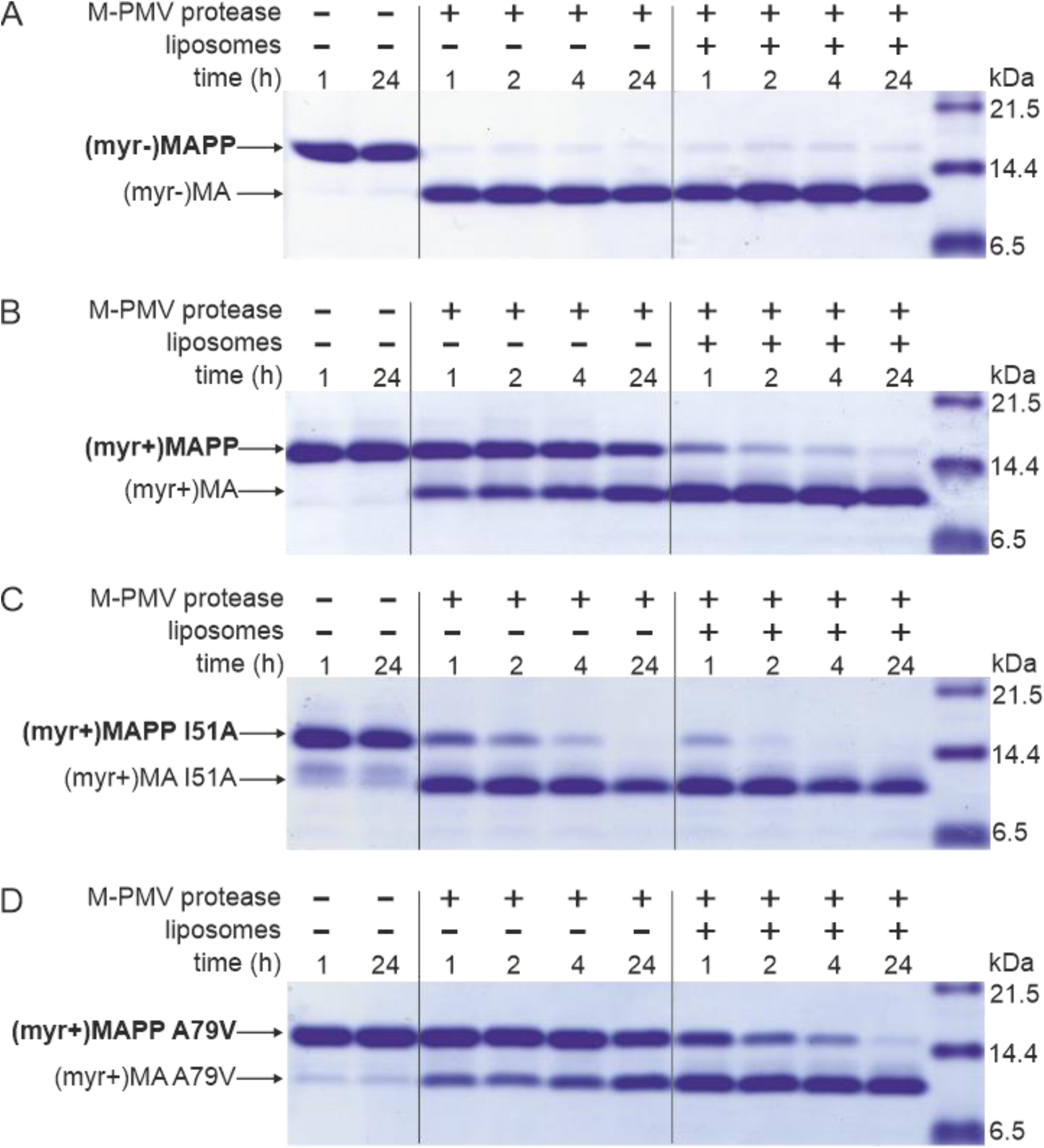
MAPP cleavage by M-PMV protease. (A) (myr-)MAPP, (B) (myr+)MAPP, (C) “myrOUT” mutant (myr+)MAPP I51A and (D) “myrIN” mutant (myr+)MAPP A79V were cleaved by M-PMV protease in the absence or presence of liposomes mimicking the PM inner leaflet.

### The affinity of myristoyl for the hydrophobic pocket of MA modulates MA cleavage from the rest of Gag

To study the effect of myristoyl switch on the accessibility of (myr+)MAPP C-terminus to the M-PMV protease, we prepared mutants that either facilitate myristoyl release (“myrOUT”) or prevent the switch (“myrIN”) by reducing or enhancing the hydrophobicity of the protein core, respectively. To design the “myrOUT” mutant, we identified large hydrophobic amino acid residue I51 which is in direct contact with the myristoyl in M-PMV (myr+)MAPP (9) and substituted it with alanine (I51A) (**Fig. S1**). For the “myrIN” mutant, we introduced the A79V mutation previously suggested to affect myristoyl accessibility in M-PMV (myr+)MA (36) (**Fig. S1**). None of these mutations lie near the cleavage site of the MA/PP junction, to exclude the possibility of its direct structural alteration. We also measured HN-HSQC spectra of both (myr+)MAPP mutant proteins and compared them with the wt spectrum. We observed bigger chemical shift changes only for the signals of residues in proximity of the mutation which proves that neither mutation has altered the overall fold of the protein (**Fig. S2**).

Here we show that “myrOUT” (myr+)MAPP mutant was cleaved more efficiently than WT (myr+)MAPP (compare **Fig. 1C and 1B**), but less efficiently than the WT (myr-)MAPP (**Fig. 1A)**, that fully mimics the “myrOUT” conformation. The effect of the “myrIN” substitution in (myr+)MAPP was apparent only upon interaction with liposomes, when preventing the switch should reduce cleavage efficiency compared to the WT (myr+)MAPP. As expected, in the absence of liposomes, both the “myrIN” mutant (**Fig. 1D)** and WT (myr+)MAPP (**Fig. 1B)** were cleaved poorly. More importantly, in the presence of liposomes, the cleavage of “myrIN” mutant was slower than that of WT (myr+)MAPP (compare **Fig. 1D and 1B**). These results show that the myristoyl switch enables cleavage of (myr+)MAPP by M-PMV protease and the sequestered myristoyl delays the cleavage.

### Liposome interaction and myristoyl switch induces (myr+)MAPP oligomerization

Based on the known structure of the M-PMV (myr-)MA trimer (Vlach et al., 2009), we designed a mutant allowing covalent cross-linking of monomers to enrich transiently formed MA trimers. We used an approach similar to that of Tedbury *et al*. for HIV-1 MA (Tedbury et al., 2016). They introduced cysteines at positions G62 and S66 of HIV-1 MA, allowing spontaneous disulfide bridge formation between MA molecules. Accordingly, we replaced threonine at position 69 of M-PMV MAPP with a cysteine residue for trimer stabilization. In the M-PMV (myr-)MA trimer, T69 is located directly opposite C62 from the neighboring MA monomer (**Fig. 2A**). Thus, its replacement with cysteine allows spontaneous formation of a disulfide bridge under non-reducing conditions and stabilizes the trimeric form of MA.

**Fig. 2.**
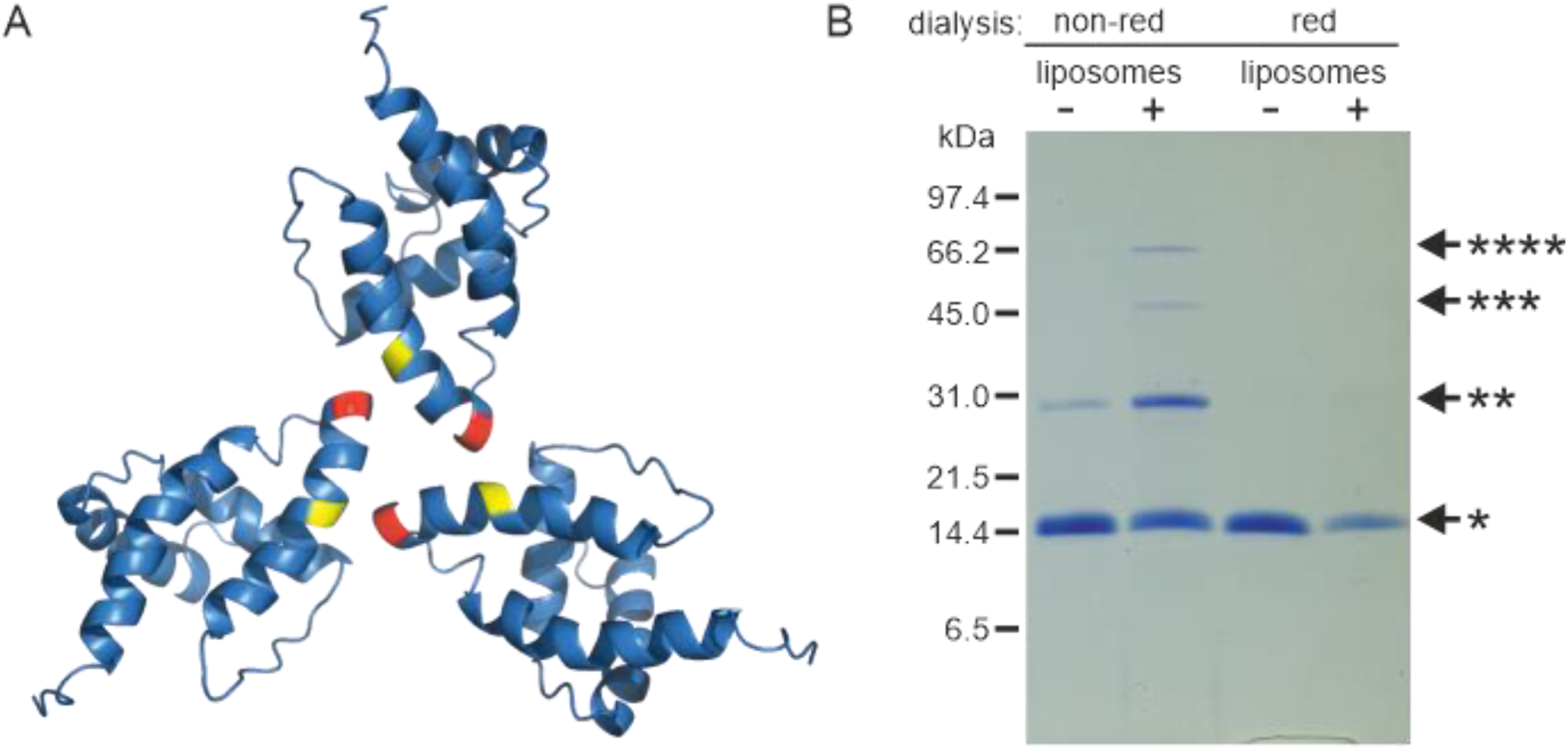
Oligomerization of (myr-)MAPP. (A) Previously published structure of WT (myr-)MA trimer (Vlach et al., 2009) with residues 62 (yellow) and 69 (red) highlighted. (B) SDS-PAGE gel showing oligomers of T69C (myr+)MAPP formed upon the interaction of T69C (myr+)MAPP with liposomes, stabilized by disulfide bridges under non-reducing conditions. non-red – non-reducing conditions; red - reducing conditions* monomer, ** dimer, *** trimer, and ****tetramer

Under reducing conditions the (myr+)MAPP T69C mutant remained monomeric both in the absence and in the presence of liposomes. Under non-reducing conditions, the (myr+)MAPP T69C mutant remained predominantly monomeric, with a small fraction of dimers. However, in the presence of liposomes under non-reducing conditions, the oligomeric fraction of the protein increased significantly, and dimers, trimers, and tetramers were formed (**Fig. 2B**). This suggests that the MA myristoyl switch that occurs after MA interaction with liposomes promotes MA oligomerization.

### Myristoyl exposure modulates the accessibility of MA junction with PP

We used hydrogen deuterium exchange (HDX)-MS to map in detail the observed differences in (myr+)MAPP and (myr-)MAPP behavior. Statistically significant differences (CI_98%_= ± 0.410 Da) were detected in regions 16-33, 67-85, and 89-110 (**Fig. S4A, Table S1A**). The first two regions of (myr-)MAPP showed considerable HDX decrease, while the 89-110 region displayed a significant increase in deuterium accessibility compared to (myr+)MAPP (**Fig. 3B)**. This shows that the absence of myristoyl in (myr-)MAPP can destabilize the alpha-helical secondary structure in the K92-L110 region, allowing it to unfold and become more flexible to facilitate proteolytic cleavage.

**Fig. 3.**
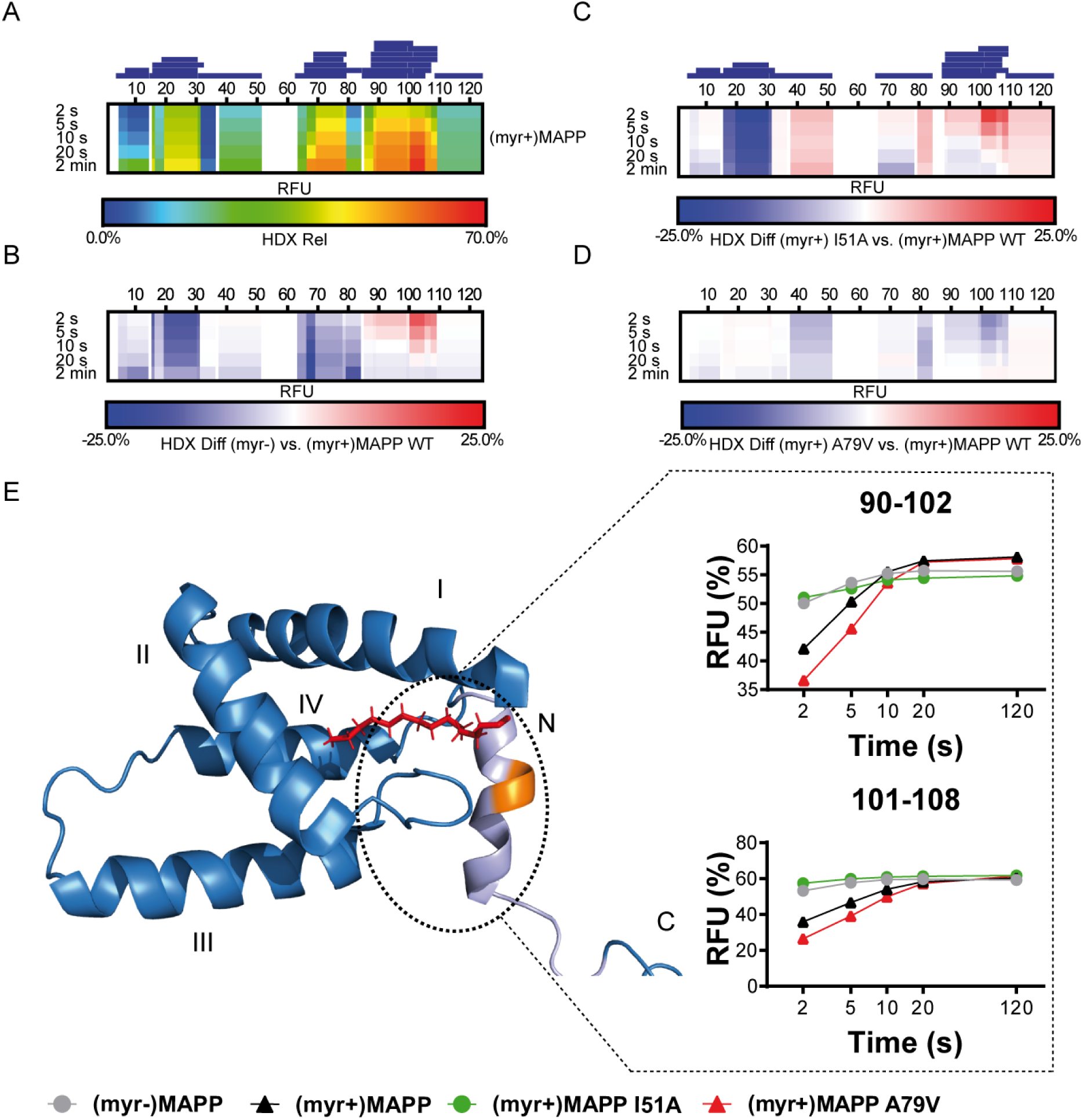
HDX of (myr+)MAPP WT and its comparison to (myr-)MAPP WT and I51A and A79V mutants of (myr+)MAPP. The heat maps were generated based on the identified peptides displayed as blue rectangles above the amino acid numbering. Each line represents one time point from the examined time course (2 s - 120 s). The regions with no HDX data are displayed as white gaps. (A) The HDX heat map of (myr+)MAPP WT shows the accessibility of each (myr+)MAPP WT amino acid in percentage of relative fractional uptake (RFU) scaled in rainbow colors. (B) Differential HDX heat map of (myr-)MAPP WT compared to (myr+)MAPP WT. ( C) Differential HDX heat map of (myr+)MAPP I51A compared to (myr+)MAPP WT. (D) Differential HDX heat map of (myr+)MAPP A79V compared to (myr+)MAPP WT. Less accessible regions are displayed in blue, while more accessible ones are in red. (E) The previously published structure of (myr+)MAPP (Prchal et al., 2012, RCSB PDB: 5LMY) with protease cleavage site shown in orange, myristoyl in red, residues 96-110 in light violet and first four helices of MA are numbered (corresponding to NMR observations).

The differences in dynamics of region 90-108 surrounding the protease cleavage site across the MAPP protein forms are indicated by uptake plots of peptides 90-102 and 101-108, respectively. The data are shown as the percentage of RFU for each MAPP protein form.

### Mutation that induces the myristoyl switch increases the dynamics of the (myr+)MAPP structure

To confirm the effect of the myristoyl switch, we monitored differences between deuterium incorporation into the “myrOUT” mutant I51A and WT (myr+)MAPP (**Fig. S4B**). The I51A amino acid substitution induced a structural transition from (myr+)MAPP to (myr-)MAPP-like (**Table S1B**). Specifically, we observed changes in region 89-110 (**Fig. 3C**), well indicated by peptides 90-102 and 101-108, respectively (**Fig. 3E**). The peptides 90-102 and 101-108 became more exposed in the I51A (myr+)MAPP mutant compared to WT (myr+)MAPP. The effect of the mutation strongly resembles WT (myr-)MAPP in deuteration rate, as I51A (myr+)MAPP reaches similar values through the whole monitored time course **(Fig. 3E**).

### A mutation that prevents the myristoyl switch stabilizes the (myr+)MAPP core and alpha helix in the protease cleavage site

Using HDX-MS, we observed changes in the structural dynamics of the “myrIN” mutant A79V (myr+)MAPP compared to WT (myr+)MAPP (**Fig. 3D; Fig. S4C; Table S1C**). The A79V mutation considerably stabilized the protein core. The region 89-110, which harbors the protease cleavage site, initially (after 2 s) exhibited even lower HDX levels in the A79V mutant compared to WT. A gradual increase in deuteration with a plateau reached after 20 s (**Fig. 3E, peptide 90-102**) or beyond 120 s (**Fig. 3E**, **peptide 101-108**) for both WT and the A79V (myr+)MAPP mutant reflects slower HDX kinetics caused by secondary structure present in the region comprising the protease cleavage site. In contrast, WT (myr-)MAPP and the I51A (myr+)MAPP mutant reached near-maximal deuteration levels at the initial time point (2 s) and maintained an almost-constant level during the measured time period (**Fig. 3E, peptides 90-102 and 101-108**). This suggests the absence of structure in their 89-110 region.

### A myristoyl switch modulates the secondary structure of the protease cleavage site between the M-PMV Gag MA and PP domains

Analysis of the previously reported structure (Prchal et al., 2012) of (myr+)MAPP identified an alpha helix spanning residues 98-106 that directly interacts with the myristoyl moiety. This region contains the M-PMV protease cleavage site located between residues 100 and 101 at the boundary of MA and PP (**Fig. 4A**). Direct comparison with the structure of non-myristoylated MAPP [(myr-)MAPP], which structurally mimics the protein with exposed myristoyl, was not possible due to the non-specific oligomerization of (myr-)MAPP, that lead to the broadening of the signals from the protein central part. However, we were able to assign the chemical shifts of backbone residues 95-119 in (myr-)MAPP and analyzed them using TALOS+ software (Shen et al., 2009) to predict the secondary structure of this region (**Table S2A**). This analysis showed that this entire region is unstructured in (myr-)MAPP (**Fig. 4B, Table S2B**). The same analysis of (myr+)MAPP predicted an alpha-helical region between residues 98-103. This indicates that the presence of the C-terminal fifth helix in M-PMV MA depends on sequestration of the N-terminal myristoyl in the hydrophobic pocket of the protein.

**Fig. 4.**
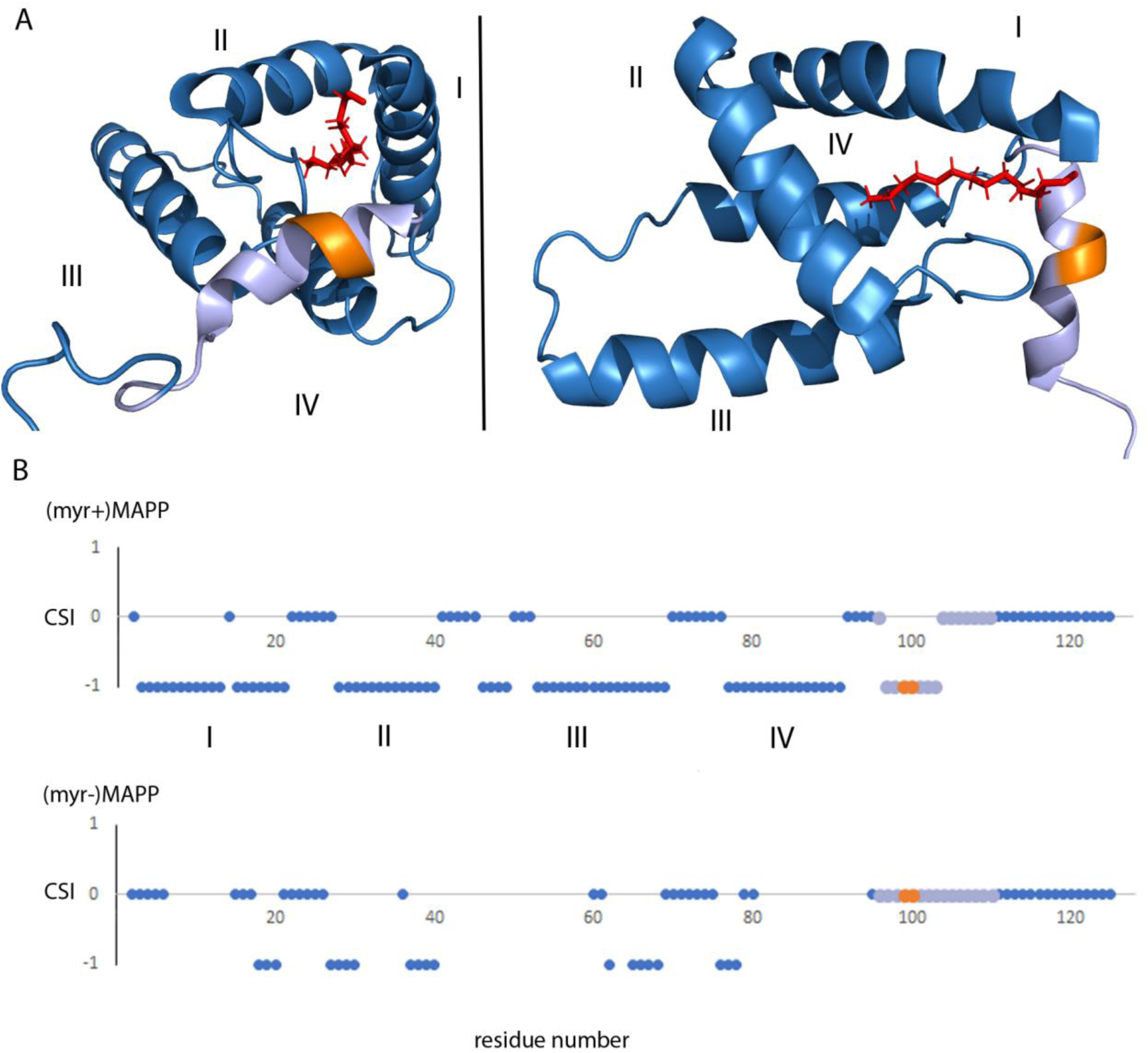
Structure of (myr+)MAPP (Prchal et al., 2012), position and secondary structure of the cleavage site between MA and PP. (A) The protease cleavage site in (myr+)MAPP is shown in orange, myristoyl in red, residues 96-110 in light violet. The first four helices of MA (blue) are numbered, the myristoyl is shown in red. (B) Backbone chemical shifts analysis of (myr+)MAPP and (myr-)MAPP proteins using TALOS+ software. CSI 0 indicate that residue is in the loop region, CSI −1 indicate that residue is in alpha-helical region. The protease cleavage site is shown in orange, residues 96-110 in light violet. The first four helices of MA (blue) are numbered.

## Discussion

Here, we examined the hypothesis that myristoyl exposure from the hydrophobic pocket of M-PMV MA triggers a conformational change at the C-terminus, facilitating proteolytic cleavage at the MA/PP junction in the Gag polyprotein. This hypothesis was based on previous results showing that non-myristoylated MA, in contrast to the myristoylated version, is efficiently cleaved from its downstream sequence in M-PMV Gag (Prchal et al., 2011). Our basic assumption was that myristoylated M-PMV MA achieves a conformation similar to that of non-myristoylated MA upon the exposure of myristoyl from the MA during the interaction with the PM.

There is a logical basis for proteolytic release of MA from M-PMV Gag to be driven by a myristoyl switch at PM. In D-type retroviruses, pre-assembled ICAPs consisting of N-terminally myristoylated Gag must travel to the PM. Premature intracytoplasmic cleavage of (myr+)MA from Gag would prevent transport of ICAPs to the PM. This was documented by Rhee and Hunter, who showed that mutant non-myristoylated M-PMV particles failed to reach the PM and accumulated in the cytoplasm (Rhee & Hunter, 1987).

Unlike in HIV-1, in M-PMV the MA myristoyl switch has not been proved. In HIV-1, the myr exposure is triggered by the interaction of the MA with PI(4,5)P_2_ (Ono et al., 2004; Saad et al., 2006; Tang et al., 2004) and the equilibrium between exposed and sequestered myr states in HIV-1 MA is also concentration-dependent (Saad et al., 2008). However M-PMV MA has significantly lower affinity for water-soluble PI(4,5)P_2_ and the interaction with this PM component fails to induce the myristoyl switch in purified M-PMV (myr+)MA *in vitro* (Kroupa et al., 2016; Prchal et al., 2012).

To prove the myristoyl switch in M-PMV MA and show its possible role in the proteolytic cleavage at MA/PP junction *in vitro*, we tried to simulate the *in vivo* situation, by interacting M-PMV MAPP with liposomes mimicking the composition of an inner leaflet of the PM (Doktorova et al., 2017). And indeed, we have shown that the interaction of M-PMV (myr+)MAPP with liposomes enabled efficient cleavage of M-PMV (myr+)MA from the downstream PP domain of Gag, probably due to the induced myristoyl switch in (myr+)MAPP. The role of the myristoyl switch itself, rather than the overall interaction of (myr+)MAPP with liposomes, on (myr+)MAPP cleavage was demonstrated using the “myrOUT” mutant, where the isoleucine residue at position 51, which is in direct contact with myristoyl in the M-PMV (myr+)MAPP (Prchal et al., 2012), was substituted by alanine (I51A). This amino acid substitution should facilitate myristoyl release by reducing the hydrophobicity of the protein core. The “myrOUT” (myr+)MAPP I51A mutant was cleaved more efficiently than WT (myr+)MAPP, but less efficiently than the WT (myr-)MAPP, that fully mimics the “myrOUT” conformation, confirming the impact of the myristoyl switch on the cleavage. Furthermore, the role of interaction of (myr+)MAPP with liposomes in (myr+)MAPP myristoyl switch was confirmed using the “myrIN” mutant, which carries the A79V substitution known to impact myristoyl accessibility in M-PMV (myr+)MA (Conte et al., 1997). In the presence of liposomes, the was cleaved more slowly than the WT (myr+)MAPP. This indicates that the interaction of (myr+)MAPP with liposomes activates the myristoyl switch in (myr+)MAPP, allowing for subsequent cleavage.

We used HDX-MS to confirm differences in accessibility of the MA/PP junction in (myr+)MAPP and (myr-)MAPP. These data showed that the absence of the N-terminal myristoyl in the binding pocket destabilizes the otherwise alpha-helical secondary structure in the K92-L110 region of (myr+)MAPP (Prchal et al., 2012), which makes the protease cleavage site more accessible. The effect of the myristoyl switch was further confirmed by analysis of cleavage of (myr+)MAPP protein carrying amino acid substitution I51A. This “myrOUT” mutant was cleaved more efficiently than WT(myr+)MAPP. HDX-MS analysis documented changes in the region spanning residues 89-110 (**Fig. 3B-D**), most prominent in the peptides 90-102 and 101-108 (**Fig. 3E**). In contrast, the “myrIN” A79V (myr+)MAPP mutant with tightly sequestered myristoyl group was cleaved less efficiently in the presence of liposomes compared to WT. The tightly sequestered myristoyl hindered the myristoyl switch in liposome-bound “myrIN” mutant, slowing the cleavage. Results of HDX-MS analysis also showed a similar pattern of 89-110 region deuteration kinetics for the “myrIN” mutant and WT (myr+)MAPP (**Fig. 3D**). In both the “myrIN” mutant and WT (myr+)MAPP, the plateau of deuteration for peptides 90-102 and 101-108 was achieved only after an extended period of 20 s (**Fig. 3E, peptide 90-102**), or after 120 s. (**Fig. 3E, peptide 101-108**). Similarly, the well-cleaved “myrOUT” mutant was almost deuterium-saturated after only 2 s and showed no significant evolution after that, suggesting that the region spanned by peptides 90-102 and 101-108 is dynamic and highly accessible for deuterium exchange (**Fig. 3E)**. The fast HDX kinetics of “myrOUT” mutant corresponds with their higher susceptibility to proteolytic cleavage compared to WT (myr+)MAPP and “myrIN” mutant. The differences in dynamics reflecting protease cleavage site accessibility could be monitored for up to 5 s with a dramatic difference in the first 2 s. These results support our data suggesting that the sequestered myristoyl participates in the mechanism preventing proteolytic separation of MA from the rest of Gag prior to its interaction with the PM.

Analysis of the secondary structure parameter estimation suggested that the region spanning amino acid residues 98–103 in (myr-)MAPP is unstructured and lacks any periodic secondary structure. In contrast, the region spanning these residues in (myr+)MAPP is ordered into the terminal fifth alpha helix of MA (Prchal et al., 2012). Residue 98 directly interacts with the myristoyl moiety, and residues 101 and 102 with the loop connecting helices II and III. The conformation of this loop differs depending on the myristoylation state of MA, as the loop is directly involved in the formation of the myristoyl-holding cavity. Thus, the switch can lead to a large conformational change in this region and destabilize the helix, resulting in exposure of the MA/PP cleavage site. Overall, this mechanism may prevent premature cleavage of (myr+)MA from the downstream portion of Gag during the transport of immature particles to the PM. The interaction of MA domain of M-PMV Gag with the PM triggering the myristoyl switch may be an important regulatory element of the Gag processing of D-type retroviruses.

In C-type retroviruses including HIV, myristoylation of the MA domain of Gag promotes particle assembly at the PM. This initiates the final steps of the virus life cycle: budding, protease activation, and maturation (Hermida-Matsumoto & Resh, 1999; Spearman et al., 1997). A myristoylation defect prevents HIV-1 Gag polyproteins from PM binding and protease activation in T cells (Lee et al., 1998). Recent findings suggest that HIV-1 protease may be activated prior to the release of virions (Tabler et al., 2022). In M-PMV, intracytoplasmic activation of protease was previously documented as the presence of mature proteins in transfected cell lysates (Rhee & Hunter, 1990). However, it is unclear whether this occurred in the cytosol or upon the interaction of immature particles with the PM and MA myristoyl switch.

*In vitro* experiments showed that the first cleavage in HIV-1 Gag takes place at the SP1/NC junction (Pettit et al., 1998). Next, the MA/CA and SP2/p6 cleavage sites are processed. The final cleavages occur at the CA/SP1 and NC/SP2 sites. Unfortunately, there is no detailed data on the kinetics of M-PMV Gag processing. However, *in vitro* experiments with immature particles isolated from COS-1 cells indicated that 120 min after DTT-induced activation of M-PMV protease, the Gag cleavage products MA, PP, and CA were present at 40%, 35%, and 15%, respectively, compared to the DTT-untreated control (Parker & Hunter, 2001). Parker and Hunter (Parker & Hunter, 2001) observed very slow release of mature M-PMV (myr+)MA from the downstream Gag region, however, available data indicated that the M-PMV MA/PP junction is cleaved more readily than other processing sites. Notably, these HIV-1 and M-PMV *in vitro* experiments using myristoylated proteins occurred in the absence of membranes. Under these conditions, Gag molecules were in a myristoyl-sequestered form.

In addition to aiding Gag processing, myristoyl exposure at the PM modulated the oligomerization of M-PMV MA. Although tertiary structures of MA proteins are very similar across retroviruses, they vary in their primary structures and surface features, including the surface distribution of hydrophilic and hydrophobic residues. In HIV-1, (myr+)MA occurs in a momomer-trimer equilibrium, with the myristoyl sequestered in the monomer and exposed in the trimer (Tang et al., 2004). Moreover, MA trimers in HIV-1 particles have been observed *in vivo* (Tedbury et al., 2016). Interestingly, structurally similar HIV-2 MA is monomeric in both its myristoylated and non-myristoylated forms (Saad et al., 2008). Previously, we showed that, unlike HIV-1 (myr+)MA, M-PMV (myr+)MAPP does not oligomerize *in vitro* (Prchal et al., 2012) and (myr-)MA exists in a monomer-dimer-trimer equilibrium (Srb et al., 2011; Vlach et al., 2009). These findings along with our cleavage experiments, justifies the hypothesis that the myristoyl switch converting M-PMV (myr+)MA into (myr-)MA-like conformation promotes its oligomerization. This suggests that the presence of the myristoyl in the hydrophobic cavity of (myr+)MA affects the hydrophobic interactions occurring at the oligomerization interfaces of the monomers. This was confirmed by using the T69C mutant with stabilized (myr+)MAPP oligomers through disulfide bridges. In the absence of liposomes, we observed only the monomeric fraction, but in their presence, we also observed dimers, trimers, and even tetramers (**Fig. 2B**). Our previous research indicated that (myr-)MA at the same concentration primarily forms dimers and some trimers (Prchal et al., 2012). In the T69C trimer, C69 is located directly opposite C62 from the second monomer (**Fig. 2A**). Dimers are likely stabilized by disulfide bridges between C42 residues (**Fig. S3**). We also observed tetramers, which we believe are two non-specifically cross-linked dimers.

Our findings reveal that the PM interaction induces myristoyl exposure from the hydrophobic core of M-PMV MA. The myristoyl switch then supports both proteolytic separation of MA from the downstream part of Gag and MA oligomerization. Broadly, these results suggest that similar mechanisms of functional modulation may occur in domains of other myristoylated proteins. For example, recoverin and calcium- and integrin-binding protein 2 exploit a myristoyl switch to modulate calcium binding (Ames et al., 1997).Thus, myristoyl exposure-triggered structural changes may have more general validity and may play various regulatory roles in other N-terminally myristoylated proteins.

## Materials and methods

### Vectors

All mutations in the vector pET22bMAPPHis (Prchal et al., 2011) were introduced using Efficient Mutagenesis Independent of Ligation (EMILI) (Fuzik et al., 2014). The mutation responsible for amino acid substitution I51A was introduced using primers MAPPHisI51Af GGAACCGCAGATATTAAACGGTGGCGTAGAG and MAPPHisI51Ar CGTTTAATATCTGCGGTTCCCTCTTGCGG, for the amino acid substitution A79V using primers MAPPHisA79Vf TAACTGTTTTCTCTTACTGGAACTTAATTAAAGAATTGATAGATAAG and MAPPHisA79Vr TAAGAGAAAACAGTTACTGGGACTTTCTCCG, and for amino acid substitution T69C using primers MAPPHisT69Cf TATTACAATTGTTTTGGCCCGGAGAAAGTCCC and MAPPHisT69Cr GGGCCAAAACAATTGTAATAGTCTTGGAAACAGTCG.

### Recombinant protein production and purification

All (myr-) and (myr+)MAPPs were produced in *E. coli* BL21 (DE3) cultivated in LB medium and purified using metal affinity chromatography on Ni-NTA agarose according to a previously published protocol (Prchal et al., 2011). The (myr+)MAPP T69C protein was purified under reducing conditions. Uniformly isotopically labeled proteins were produced using M9 minimal medium (Hiatt et al., 2018) with [U–^15^N]NH4Cl and D–[U–^13^C]glucose (CIL, USA). The identities of the prepared proteins and the degrees of myristoylation were confirmed using MALDI-TOF/TOF mass spectrometry on an Autoflex speed mass spectrometer (Bruker Daltonics). The 13 kDa form of M-PMV protease was prepared using a previously published protocol (Zabransky et al., 1998).

### Liposome preparation

To prepare liposomes mimicking the PM inner leaflet (Doktorova et al., 2017), individual lipids (purchased from Avanti Polar Lipids, Inc.) were dissolved in chloroform or in chloroform/methanol/water (20:9:1) in the case of PI(4,5)P_2_ and thoroughly mixed to a final lipid concentration of 5 mg/ml. To obtain 250 µl of lipid mixture, we mixed 310 µg of cholesterol, 400 µg of phosphatidylethanolamine, 75 µg of phosphatidylcholine, 290 µg of phosphatidylserine, 38 µg of PI(4,5)P_2_ and 140 µg of phosphatidylinositol. Chloroform was evaporated, and the lipid mixture was resuspended in a protease cleavage buffer or in phosphate buffered saline (PBS). Liposomes were formed using a mini-extruder (Avanti Polar Lipids, Inc.) with a 100-nm polycarbonate filter.

### MAPP cleavage by M-PMV protease

All (myr-) and (myr+)MAPP proteins were cleaved by M-PMV protease in the protease cleavage buffer (50 mM acetate, pH 5.3, 300 mM NaCl, 0.05% mercaptoethanol). Briefly, 80 µg of each protein solution was incubated with 2U of protease (1U of M-PMV protease cleaves 100 µg of (myr-)MAPP in 1 h) in a total volume of 220 µl of protease cleavage buffer in the absence or presence of 20 µl liposomes. Aliquots (20 μl each) were collected at time intervals of 1 h, 2 h, 4 h and 24 h, resuspended in 2x reducing protein loading buffer (PLB), and analyzed by Tris-Tricine SDS PAGE.

### Interaction of (myr+)MAPP T69C with liposomes

A 40 µg portion of (myr+)MAPP T69C solution in reducing PBS was mixed with liposome suspension in a 1:1 (v/v) ratio to obtain a total volume of 100 µl. Protein diluted 1:1 with reducing PBS was used as a control. The samples were dialyzed overnight either against non-reducing or reducing PBS. Aliquots were collected, resuspended in 2x non-reducing PLB, and analyzed by Tris-Tricine SDS PAGE.

### Hydrogen-deuterium exchange

For HDX labeling experiments, 0.2 mM wild type (WT) (myr-)MAPP, WT (myr+)MAPP, and the I51A, I86A, A79L, and A79V mutants were mixed with D_2_O in a 1:9 ratio. Samples were incubated at 4 °C for 0, 2, 5, 10, 20 or 120 seconds and quenched with an equal volume of quench buffer (8 M urea, 1 M Glycine, pH 2.51). In case of the (myr+)MAPP I51A mutant, 200 mM TCEP was added to the quench buffer to increase its sequence coverage. The HDX experiments were performed in triplicates for each labeling time point.

### LC-MS analysis

Peptides were identified by tandem mass spectrometry of non-deuterated proteins. Samples were injected into a refrigerated UPLC system (NanoAcquity, Waters) with chromatographic elements held at 0 °C. The samples were then passed through a pepsin protease column (Enzymate Protein Pepsin Column, 300Å, 5 µm, 2.1 mm x 30 mm, Waters) at 15 °C and flow rate of 100 µl min^-1^ (0.1% v/v FA), and the generated peptides were trapped and desalted for 3 minutes on a VanGuard pre-column (2.1 mm x 5 mm Waters Acquity UPLC BEh C_18_ (pore size 1.7µm)). Peptides were then separated in gradient of acetonitrile for 12 min over an Acquity UPLC column (1 mm x 100 mm, 1.7 µm BEH C_18_) for 12 min (10-35% CH3CN v/v and 0.1% v/v FA, flow rate 40 µl min^-1^). The MS spectra were acquired with a Synapt G2 mass spectrometer (ESI-Q/TOF; Waters) in mass range from 50 to 2000 m/z with Leu-enkephalin serving as a continuous (lock-spray) calibration standard and performing a scan every 0.4 s. The list of peptides was obtained by using ProteinLynx Global SERVER™ (PLGS; Waters) version 3.0.2 with processing parameters as follows: chromatographic peak width – automatic, MS TOF resolution – automatic, Lock Mass for Charge +1 – 556.2771 Da/e, Lock Mass Window - 0.25 Da, Low Energy Threshold – 135.0 counts, Elevated Energy Threshold – 30.0 counts and Intensity Threshold – 750.0 counts. PLGS workflow parameters were as follows: searching against fasta file containing forward and reverse sequences of examined proteins and Pepsin (Uniprot code P00791), peptide tolerance and fragment tolerance – automatic, minimum fragment ion matches per peptide – 3, minimum fragment ion matches per protein – 7, minimum peptide matches per protein – 1, primary digest reagent – non-specific, number of missed cleavages 3, oxidation of methionines as a variable modifier reagent, false discovery rate 5, monoisotopic mass of peptides with charge +1. The LC analysis of labeled samples was identical to that of non-deuterated samples.

DynamX 3.0 (Waters) was used to filter peptides for the determination of HDX differences between examined proteins by selecting those having the length up to 25 amino acids, presenting 0.3 fragment per amino acid and showing mass error for the precursor ion below 10 ppm. In addition, only the peptides that were identified in at least 3 out of 5 of the acquired MS/MS files and having a minimum signal intensity 3000 with retention time RSD up to 5 % were used for further analysis. MS Files were processed according to the parameters as follow: both chromatographic peak width and MS TOF resolution as automatic, 556.2771 Da as a Lock mass for charge +1, Lock mass window 0.25 Da, Low energy threshold 130 and elution time range to search in for the data 2.5-9 min. DynamX advanced processing parameters were not applied.

### NMR spectroscopy

All NMR data were collected on a Bruker AvanceIII 600-MHz NMR spectrometer equipped with a cryoprobe (Bruker BioSpin, GmbH, Germany). The backbone atoms of (myr+)MAPP were assigned using the standard set of triple-resonance experiments (HNCA, HN(CO)CA, HNCACB, CBCA(CO)NH and HNCO). An estimate of CSI and the backbone dihedral angles Φ and Ψ was performed with TALOS+ software and was based on the ^1^HN, ^13^CO, ^13^Cα, ^13^Cβ and ^15^NH chemical shifts. The data were processed with TopSpin (Bruker BioSpin GmbH, version 3.6) and analyzed using CcpNmr analysis (Vranken et al., 2005). Structures were visualized with the PyMOL Molecular Graphics System (Version 2.0 Schrödinger, LLC).

## Acknowledgments

The research was supported by the Czech Science Foundation (grant No. 22-19250S) and by the project National Institute of virology and bacteriology (Programme EXCELES, ID Project No. LX22NPO5103) - Funded by the European Union - Next Generation EU. We would like to thank Petra Junkova for careful reading and suggestions to improve the manuscript.

## Classification

Biochemistry and Chemical Biology, Microbiology and Infectious Disease

---

**Fig. S1.**
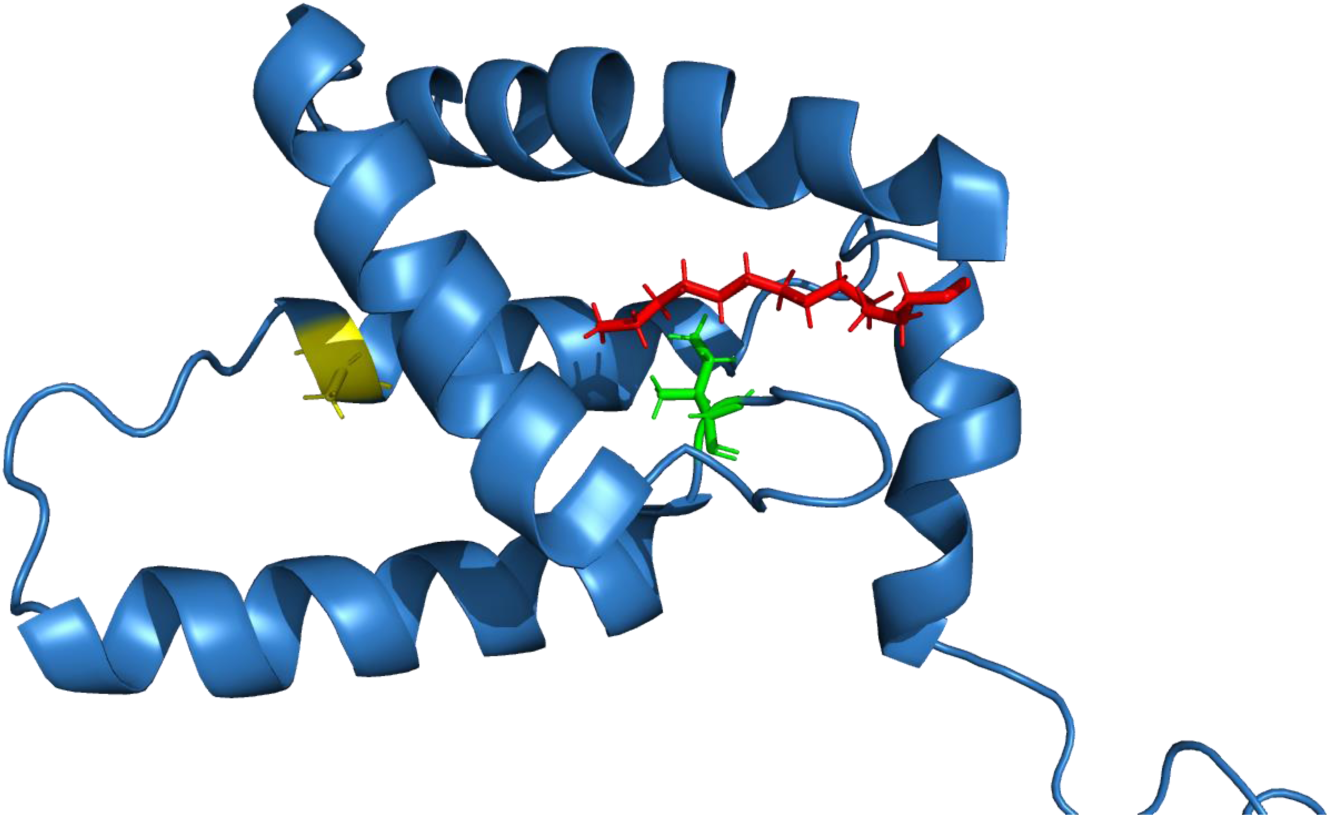
Structure of WT (myr+)MAPP indicating positions of the mutated residues The positions of mutated residues are indicated in previously published structure (Prchal et al., 2012): I51 (green) and A79 (yellow). Myristoyl is shown in red.

**Fig. S2.**
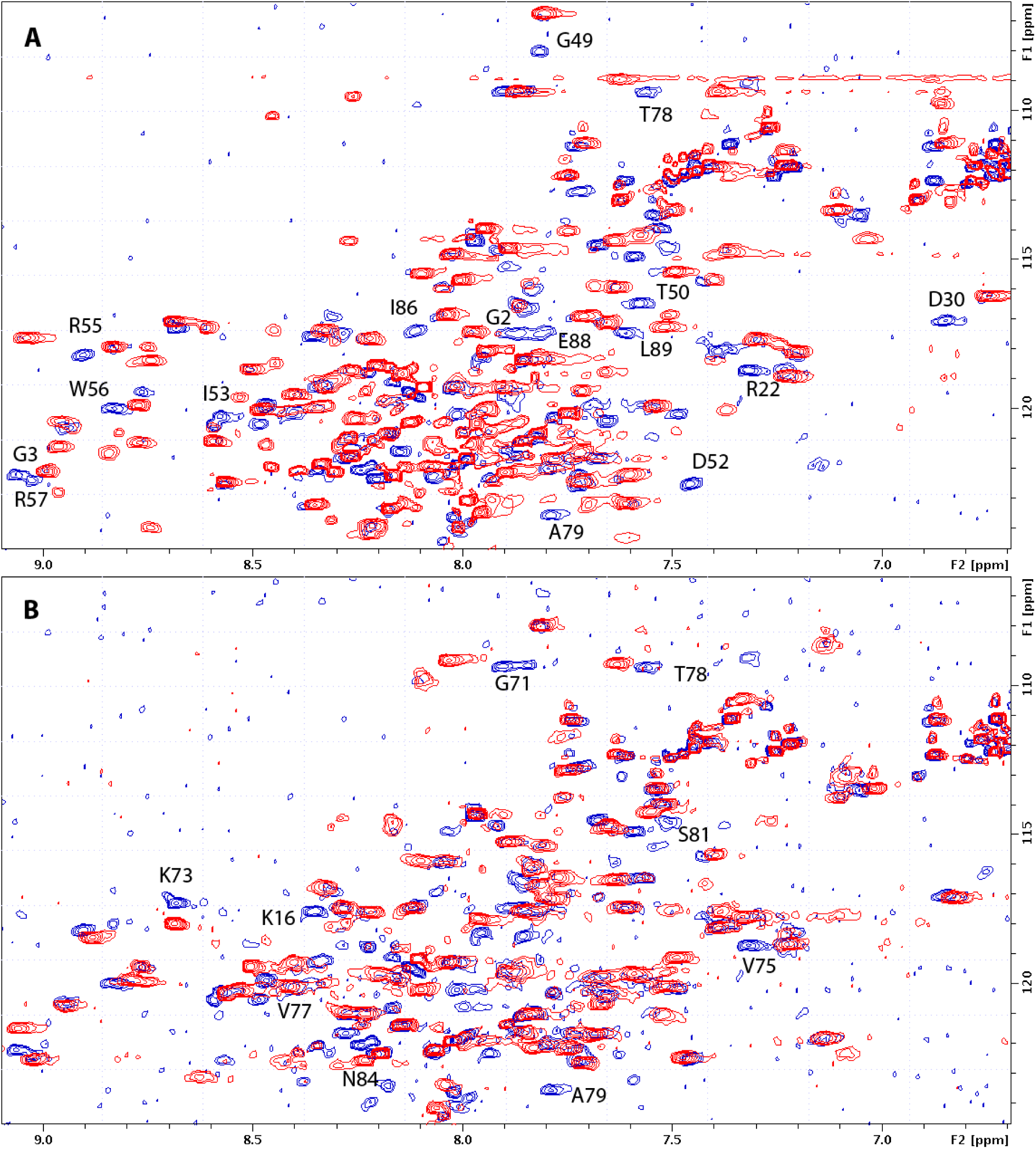
Comparison of (myr+)MAPP wt and mutants HN-HSQC spectra. HN-HSQC spectra of wt (myr+)MAPP (blue) overlaid with the same spectra of I51A (red, panel A) and A79V (red, panel B) mutants. Signals of wt HN groups that are not overlapped by signals of mutant are assigned.

**Fig. S3.**
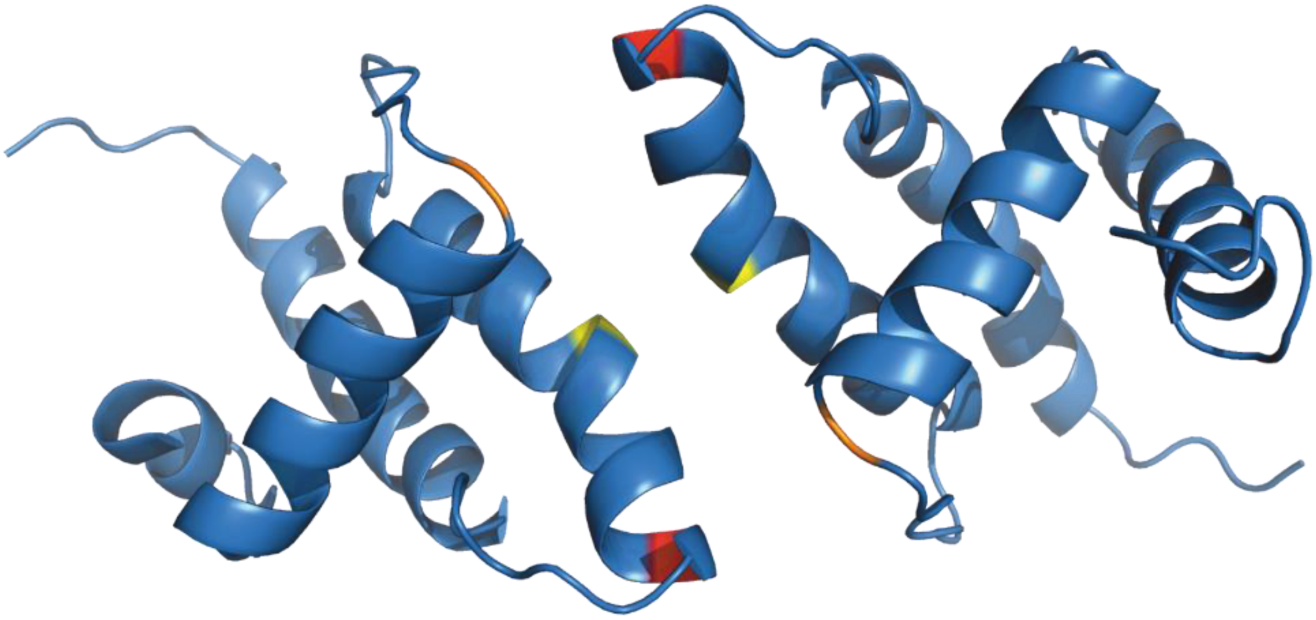
Structure of (myr-)MA wild type dimer (Vlach et al., 2009) Previously published ribbon structure of (myr-)MA wild type dimer **(Vlach et al., 2009)** showing positions of residues T69 (in red), C62 (in yelow) and C42 (in orange).

**Fig. S4A.**
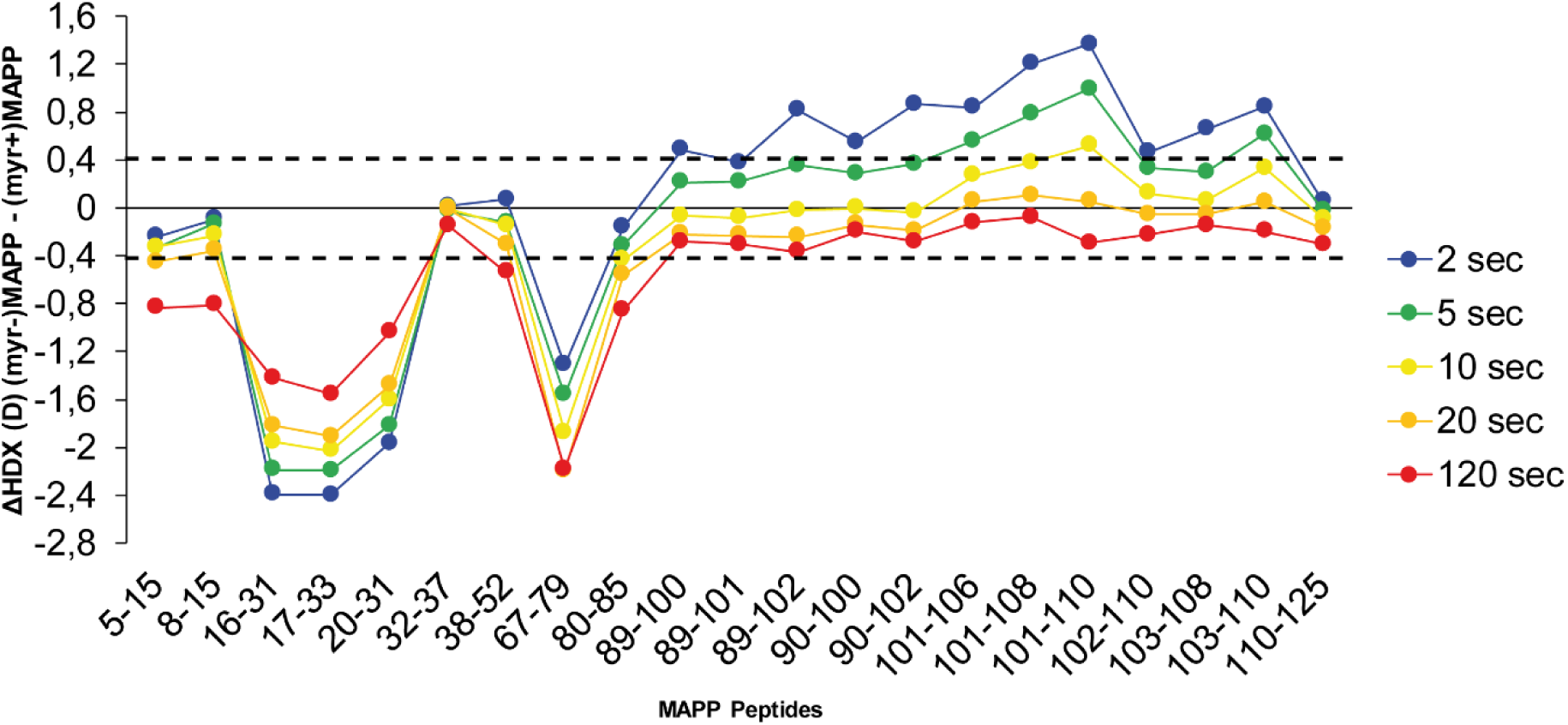
Comparison of HDX of (myr+)MAPP and (myr-)MAPP wild types. The butterfly plot shows the HDX differences between WT (myr+)MAPP and WT (myr-)MAPP over the time (see also **Table S1**). Peptides with significant differences in HDX are arranged according to their positions in MAPP. Confidence threshold (CI_98%_) is displayed as a dotted black line. The value of confidence threshold was calculated as ± 0.410 Da.

**Fig. S4B.**
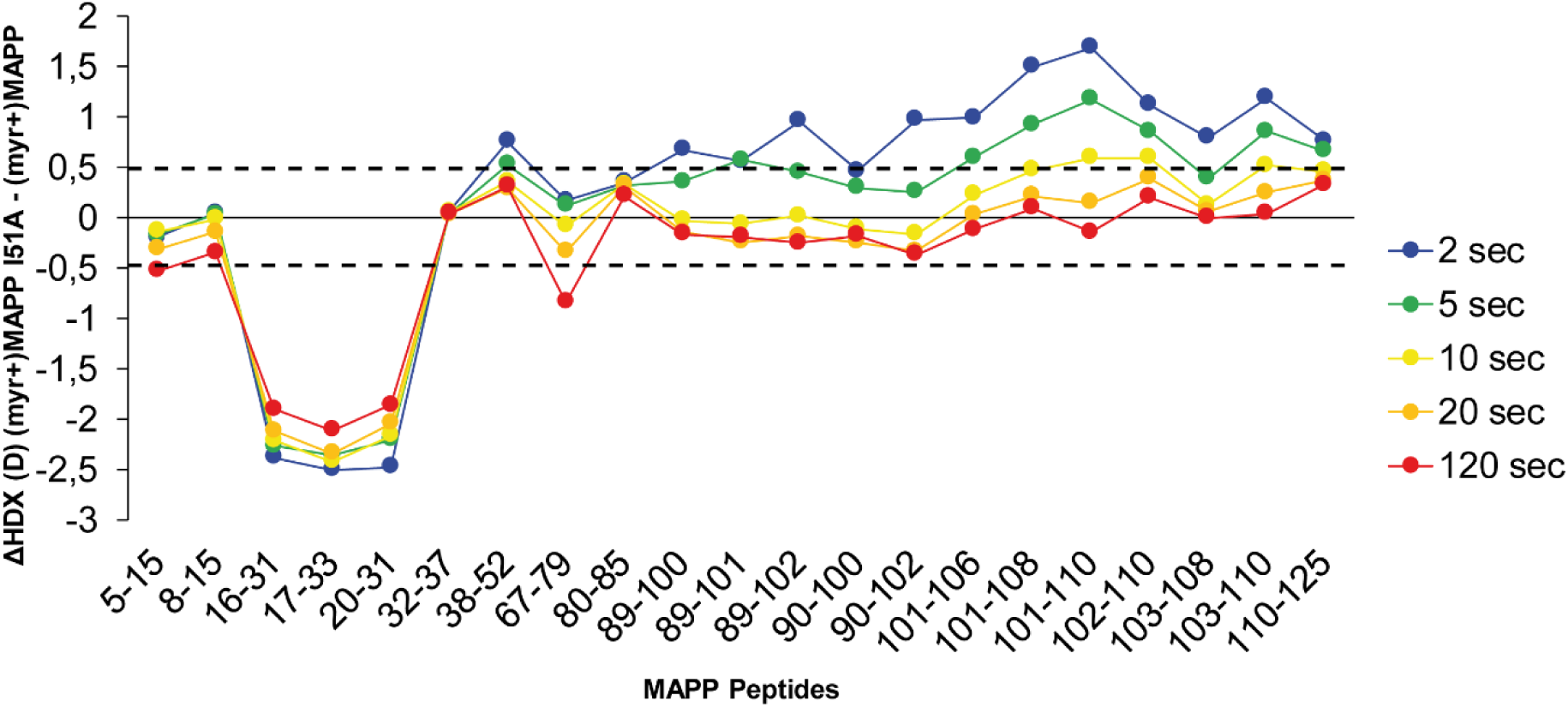
Comparison of HDX of (myr+)MAPP and I51A (myr+)MAPP. The butterfly plot shows the HDX differences between WT (myr+)MAPP and I51A (myr+)MAPP over the time (see also **Table S2**). Peptides with significant differences in HDX are arranged according to their positions in MAPP. Confidence threshold (CI_98%_) is displayed as a dotted black line. The value of confidence threshold was calculated as ± 0.478 Da.

**Fig. S4C.**
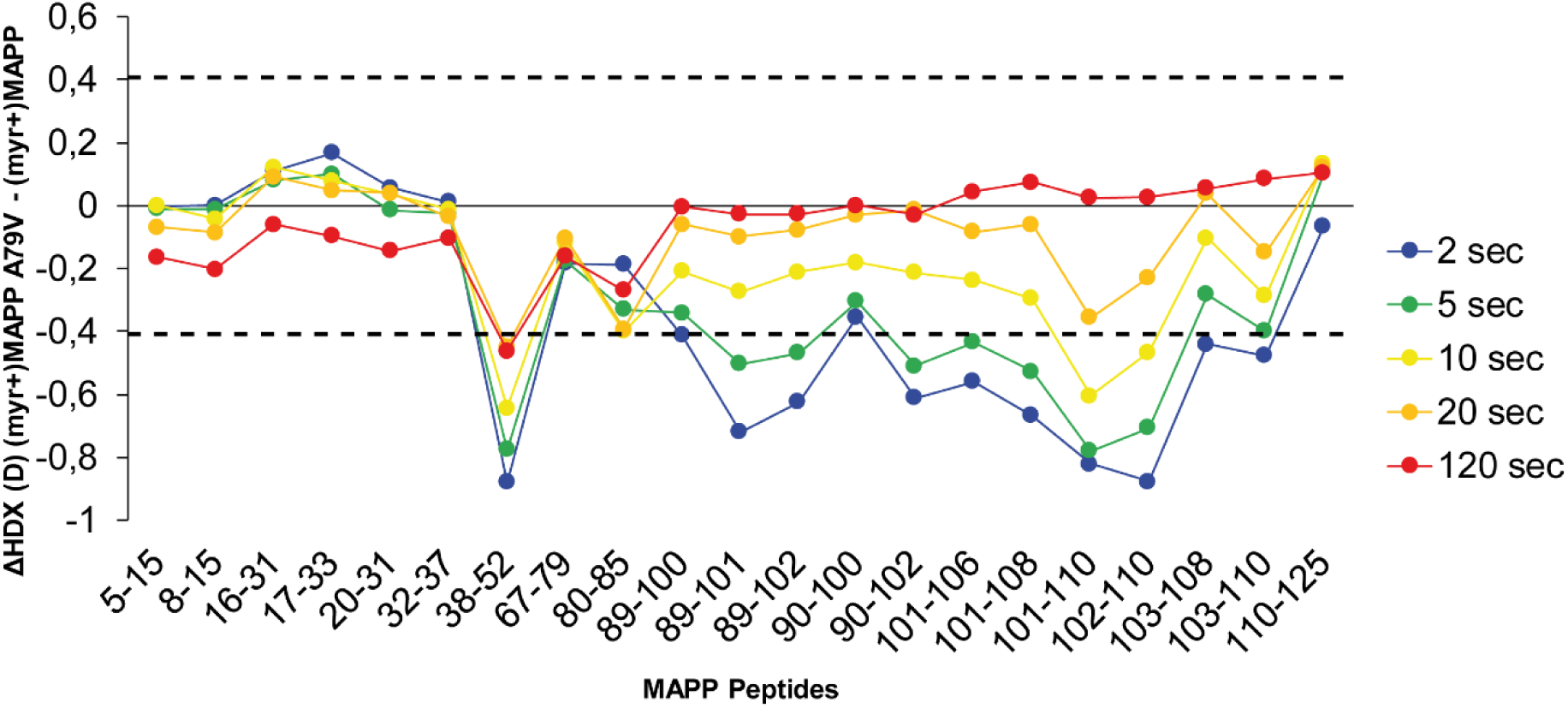
Comparison of HDX of (myr+)MAPP and A79V (myr+)MAPP. The butterfly plot shows the HDX differences between WT (myr+)MAPP and A79V (myr+)MAPP over the time (see also **Tables S3**). Peptides with significant differences in HDX are arranged according to their positions in MAPP. Confidence threshold (CI_98%_) is displayed as a dotted black line. The value of confidence threshold was calculated as ± 0.410 Da.

